# Supervising the PhD: identifying common mismatches in expectations between candidate and supervisor to improve research training outcomes

**DOI:** 10.1101/2020.02.20.958520

**Authors:** Adam P.A. Cardilini, Alice Risely, Mark F. Richardson

## Abstract

The relationship between a PhD candidate and their supervisor is influential in not only successful candidate completion, but maintaining candidate satisfaction and mental health. We quantified potential mismatches between the PhD candidates and supervisors expectations as a potential mechanism that facilitates poor candidate experiences and research training outcomes. 114 PhD candidates and 52 supervisors ranked the importance of student attributes and outcomes at the beginning and end of candidature. In relation to specific attributes, supervisors indicated the level of guidance they expected to give the candidate and candidates indicated the level of guidance they expected to receive. Candidates also report on whether different aspects of candidature influenced their mental well-being. We identified differences between candidates and supervisors perceived supervisor teaching responsibility and influences on mental well-being. Our results indicate that the majority of candidates were satisfied overall with their supervision, and find alignment of many expectations between both parties. Yet, we find that candidates have much higher expectations of achieving quantitative outcomes than supervisors. Supervisors believed they give more guidance to candidates than candidates perceive they received, and supervisors often only provided guidance when the candidate explicitly asked. Personal expectations and research progress significantly and negatively influenced over 50% of candidate’s mental well-being. Our results highlight the importance of candidates and supervisors explicitly communicating the responsibilities and expectations of the roles they play in helping candidates develop research skills. We provide four suggestions to supervisors that may be particularly effective at increasing communication, avoiding potential conflict and promoting candidate success and wellbeing.

## Introduction

Any candidate will tell you doing a PhD is hard. However, guiding a PhD candidate through the bumpy process can be just as tricky for the supervisor. Developing a good relationship between the candidate and supervisor is one of the most important components of a successful PhD. Research has consistently shown that the relationship between the candidate and their supervisor is one of the most important predictors of candidate dissatisfaction (Ives & Rowley, 2005; Kovach Clark et al., 2009; Tompkins et al., 2016), PhD discontinuation (Bair & Haworth, 2004; Buckley & Hooley 1, 1988; Golde, 2000; Kiley, 2011; Lovitts & Nelson, 2000), and depression (Peluso et al., 2011). In Australia, 20% of candidates report not being satisfied with their supervision (McGagh et al., 2016), and 20-35% are estimated to drop out of their program (Jiranek, 2010; McGagh et al., 2016). Similarly, other studies have found over 30% of PhD candidates are at risk of having or developing common psychiatric disorders (Levecque et al., 2017; Peluso et al., 2011). This equates to large-scale negative mental health and career consequences across students, as well as a significant loss of research output (Larivière, 2011). Importantly, these negative outcomes cannot be solely attributed to the candidate. Supervisor engagement, amongst other factors such as faculty support (Golde, 2000), have been increasingly linked to candidate research productivity (Gu et al., 2011), student completion rates (Buckley & Hooley 1, 1988; Kiley, 2011), and student mental health (Levecque et al., 2017; Peluso et al., 2011). As such, there has been an increasing emphasis on the importance of supervisor training in improving outcomes for candidates (Delamont et al., 2004; Halse, 2011; Halse & Bansel, 2012), as well as cognitive-behavioural coaching for candidates (Kearns, Gardiner, et al., 2008). However, although supervisor training emphasizes the importance of clear communication of expectations between both parties (Delamont et al., 2004; Moxham et al., 2013), there is currently little quantitative understanding of how common expectations are either aligned or mismatched within Australian institutions and how these can be addressed.

Mismatched expectations between candidates and supervisors have been shown to have a negative effect on candidate completion rates and timeliness(Holbrook et al., 2014; McCormack, 2004). Clear communication of expectations between candidates and supervisors is considered to be paramount to fostering a successful working relationship (Delamont et al., 2004; Moxham et al., 2013). However, expectations are challenging to address and quantify because they cover a huge array of responsibilities and outcomes that are critical to PhD completion (Mowbray & Halse, 2010), and these by necessity change over time (e.g. supervisor expectations of the student will differ between the beginning of candidature and at the time of completion). There is also an increasing distinction between expectations revolving around quantifiable student outcomes (e.g. number of papers published or grants attained) and qualitative outcomes (e.g. critical thinking or technical skills specific to the field), and these may differ between candidate and supervisor if the institution does not require a PhD by publication (Lindén et al., 2013; Vilkinas, 2008). Moreover, the extent to which the supervisor is expected to guide the candidate in developing the necessary critical skills to complete a PhD and prepare the candidate for a career in their chosen field may be another source of conflict. Critically, supervisors are increasingly time deficient and juggling many responsibilities and obligations, giving them little freedom to dedicate time to the challenges that individual candidates may face. Although the dynamics of such relationships are unique to the candidate and supervisor in question, trends in candidate and supervisor expectations may exist that allow supervisors to focus efforts on certain areas in order to promote more effective communication between the two.

In this study, we aimed to identify the importance of common expectations surrounding the candidate – supervisor relationship. By identifying expectations misaligned in candidates and supervisors, we can understand where efforts should be directed to avoid conflict and promote candidate success. 114 PhD candidates and 52 supervisors answered a questionnaire that asked them to rank common expectations and outcomes of students by importance for both the beginning and end of candidature. In addition, we also assessed the level of guidance expected to be given to the student by the supervisor for each outcome, in order to identify discrepancies between how students and supervisors perceive supervisor responsibility. We then compared answers between students and supervisors for both analyses to identify where the biggest discrepancies in expectations lay. Finally, we assessed whether supervisor guidance was perceived to be linked to student well-being, in order to assess any negative impacts of supervision on student mental health. We summarise the results by providing four suggestions to supervisors that may be particularly effective at increasing communication, avoiding conflict and promoting candidate success.

## Methods

### The survey

A survey was developed to determine the difference in PhD candidate and supervisor expectations of the candidates’ attributes and supervisor guidance (supplementary file). The first question had participants self-identify as a PhD supervisor, PhD candidate, recently graduated PhD (<2 yr) or a discontinued candidate. Candidates and supervisors were given a different survey set to complete. The questions asked in each survey set were equivalent, but the questions were written for the perspective of each group (see supplementary file). For example, both candidates and supervisors received the following equivalent question respectively, *‘Please rank the quality of the PhD supervision you received’* or *‘In your opinion, rank the quality of the PhD supervision you provide*’. Each survey set included five demographic questions and six questions regarding candidate attributes, supervisor guidance and the quality of supervision. The candidate survey set included three private questions, two relating to candidate mental well-being during candidature and one on the likelihood of pursuing a career in research academia.

Two questions were asked to determine the candidate attributes that were most important to candidates and supervisors. Participants were asked to select and rank the top five attributes of a candidate starting a PhD and the top five attributes or outcomes of a candidate at the time of thesis submission (see supplementary file for attribute and outcome options sets). Two questions were asked to determine the attributes that a supervisor has the most responsibility for helping a candidate develop during a PhD. The first supervisor responsibilities question required participants to select and rank the top three attributes that a supervisor is most responsible for helping a candidate develop during their PhD (see supplementary material for attribute options). The second question asked candidates and supervisors to indicate the level of guidance they respectively received, or provide during candidature in relation to each attribute. The level of guidance was indicated on a four point scale including, *‘none’, ‘only when asked’, ‘when seen as needed’* and *‘at every opportunity’*. To determine the mismatch between candidate and supervisor experience of PhD supervision, candidates and supervisors were asked to indicated the quality of supervision they respectively received or provided during candidature.

PhD candidates were asked two questions regarding their mental well-being during candidature. The first question asked candidates to indicate whether experiences during their candidature had negatively affected their mental well-being. Responses were indicated on a five-point scale from *strongly disagree* to *strongly agree*. The second question asked candidates to indicate how significantly along a five-point scale five different aspects of candidature negatively influenced their mental well-being. The five aspects of candidature that were asked about were, *‘supervisor relationship’, ‘research environment’, ‘research progress’, ‘personal expectations’*, and *‘supervisor expectations’*. The options for the five-point scale included, *not at all significantly* to *very significantly*. Finally, candidates and supervisors were asked to rate the quality of the supervision they had received/provided along a 5 point scale including options from *very low* to *very hig*h.

### Recruitment

An email inviting people to participate in the survey, which included a link to the survey, was sent to PhD candidates and supervisors in Science and Health faculties at Deakin University, an Australian higher education institution. Schools that were expected to have a large number of PhD candidates completing research thesis were targeted, including life sciences, medicine and psychology. The survey was opened for completion from 5^th^ August 2016 to 14^th^ November 2016.

### Analysis

Question responses where participants ranked a subset of attributes created a partially ranked dataset. Each attribute that was ranked by a participant was given a numerical value equivalent to its rank, e.g. rank 1 = 1, rank 2 = 2, etc. If participants ranked the same attribute twice in a single question the second instance of the attribute was replaced with ‘NA’. To calculate the rank of all attributes, unranked attribute values must be filled in the data matrix. We populated unfilled attribute values with the average of the mean unranked value possibilities, e.g. unranked possibilities 6-16 have a mean of 11. We used the ‘rank’ function in R statistical package ‘base’ package (R Core Team, 2017) to rank each attribute. Any tied attributes were set to the maximum number of those tied attributes. We calculated separate rank values for candidate and supervisor subsets and scaled values to allow for comparison (‘scale()’, R Core Team, 2017).

We tested whether there was a significant difference in the average rank sets between candidates and supervisors using a Kendall rank correlation, tau, from the ‘Kendall’ package in R (McLeod, 2011). Correlation p-values ≤0.05 were considered significant. After scaling, rank values could be negative or positive. To make plot comparisons easier to interpret the minimum scaled rank value was added to each rank score, this made the minimum rank value equal ‘0’ and all other values positive. The highest scaled rank value corresponds to the top ranking option.

We used Generalised Linear Models (GLM) to determine differences between the levels of guidance candidates and supervisors indicated that they received/provided. We reduced the four guidance option responses into two values for analysis; *‘none’* and *‘only when asked’* responses were grouped as passive guidance and given a value of 0, while *‘when seen as needed’* and *‘at every opportunity’* were grouped as active guidance and given a value of 1. This created a binary dataset for analysis. The GLM included the binary guidance response as the response variable with candidate/supervisor status as the predictor variable. The model specified a binomial distribution.

We used Kendall non-parametric tests (‘cor.test’, R Core Team, 2017) to determine correlations between candidates reported supervisor quality, with experiences and impacts of negative well-being and career aspirations. Tau and p-values are reported. We used Wilcox non-parametric tests (‘wilcox.test’, R Core Team, 2017) to analyse the gender differences in reported supervision quality, well-being measures and career aspirations.

p-values were FDR corrected for each set of tests (Benjamini & Hochberg, 1995) and FDR corrected p-values < 0.05 were considered significant.

## Results

166 academics answered the survey on PhD expectations. Of these, 55% were current PhD candidates, 31% were supervisors, and 13% were recent graduates, while one person was a discontinued candidate (Table 1). We found a significant difference between how candidates and supervisors ranked the importance of attributes in all three of the following tests; 1) the rank of attributes candidates are expected to have at the beginning, and 2) the end of candidature, and 3) the rank of attributes supervisors are expected to provide guidance on (Table 2).

**Table 1.** Participant details

| Group | Type | N | Gender<br>F/M (Unspecified) | Age (range) |
| --- | --- | --- | --- | --- |
| Candidate | Current | 92 | 55 / 35 (1) | 32 ± 8 (22-65) |
|  | Graduate | 21 | 10 / 11 | 28 ± 10 (27-61) |
|  | Discontinued | 1 | 1 | - |
| Supervisor |  | 52 | 27 / 25 | 46 ± 9 (31 – 66) |

**Table 2.** Kendall rank correlation between the averaged attribute ranks of candidate and supervisor.

| **Rank test responses: *candidate vs. supervisor*** | **tau** | **p-value** |
| --- | --- | --- |
| Average ranked attributes of new candidates | 0.660 | <0.001 |
| Average ranked attributes of end candidates | 0.623 | <0.001 |
| Average ranked responsibility to provide guidance for attribute | 0.500 | 0.008 |

### Beginning of candidature

Expectations of both candidates and supervisors were fairly well aligned at the beginning of candidature, with both groups placing high importance on candidate motivation, enthusiasm, and written communication (Fig. 1a). However, candidates placed much higher importance on good academic grades than supervisors, whilst supervisors placed much higher importance on the candidate’s ability to think critically than did the candidates themselves. Previous publications, industry experience, self-confidence, and good verbal communication were all more important for candidates than they were to supervisors at the beginning of candidature.

**Figure 1.**
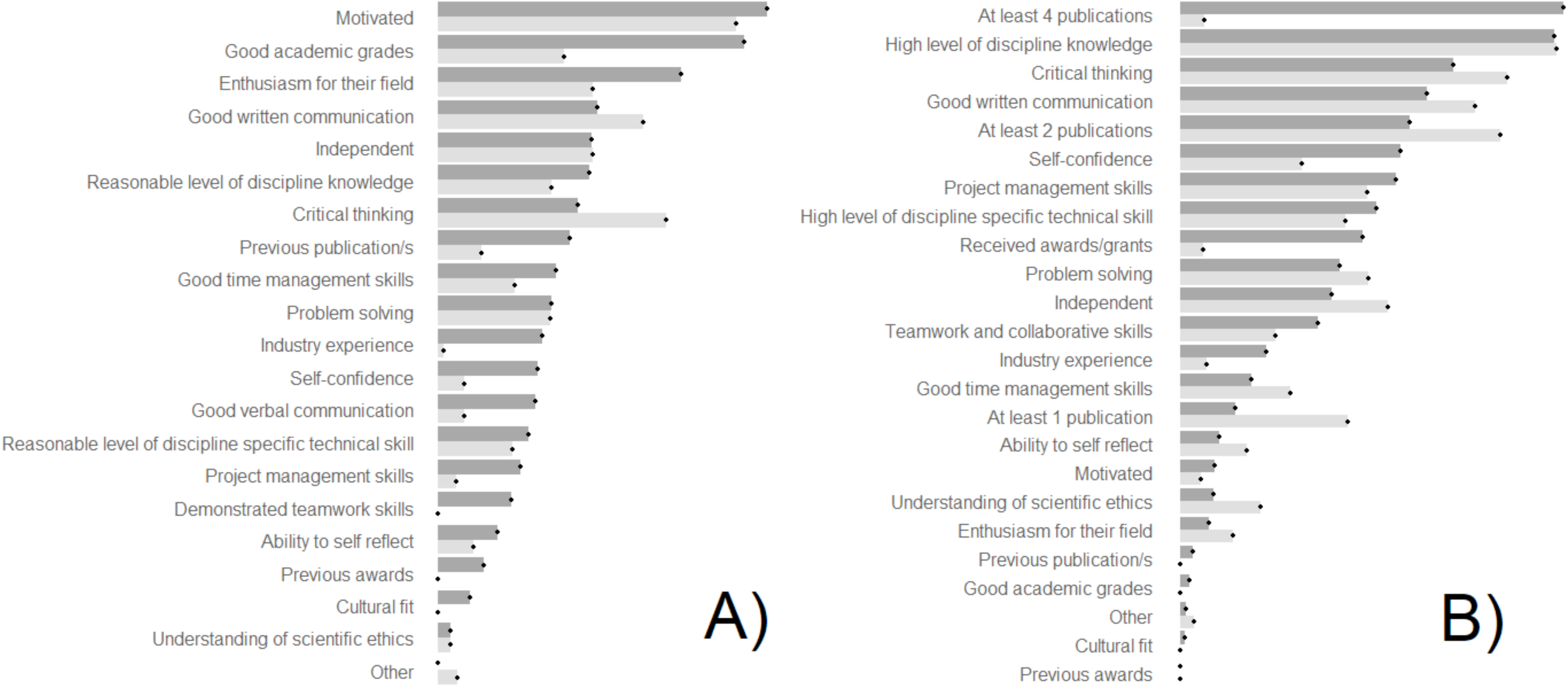
Attribute ranks as indicated by candidates (dark grey) and supervisors (light grey). Panel A) and B) show the rank of the most important attributes for a cadidate starting a PhD and finishing a PhD respectively. The relative length of the attribute bars relates to the scaled rank value that the attribute received in relation to other attributes.

### End of candidature

Overall, candidates had much higher expectations of achieving quantitative outcomes by the end of candidature than supervisors, with candidates placing high importance on publishing at least four papers, and fairly high importance on winning awards and grants (Fig. 1b). In contrast, supervisors expected the candidate to publish just one or two articles by the end of candidature, and placed low importance on winning awards and grants. Both groups placed high importance on discipline knowledge, critical thinking skills and written communication, although supervisors placed all these qualitative outcomes higher in importance than the candidates themselves.

### Responsibility to provide guidance

Both candidates and supervisors considered that supervisors have an important responsibility to provide guidance developing the candidate’s written communication skills, critical thinking skills and discipline knowledge. But candidates place the highest importance on developing written communication skills, while supervisors on critical thinking (Fig. 2). However, candidates considered that supervisors should have a slightly stronger role in encouraging or developing their academic independence, motivation and teamwork skills.

**Figure 2.**
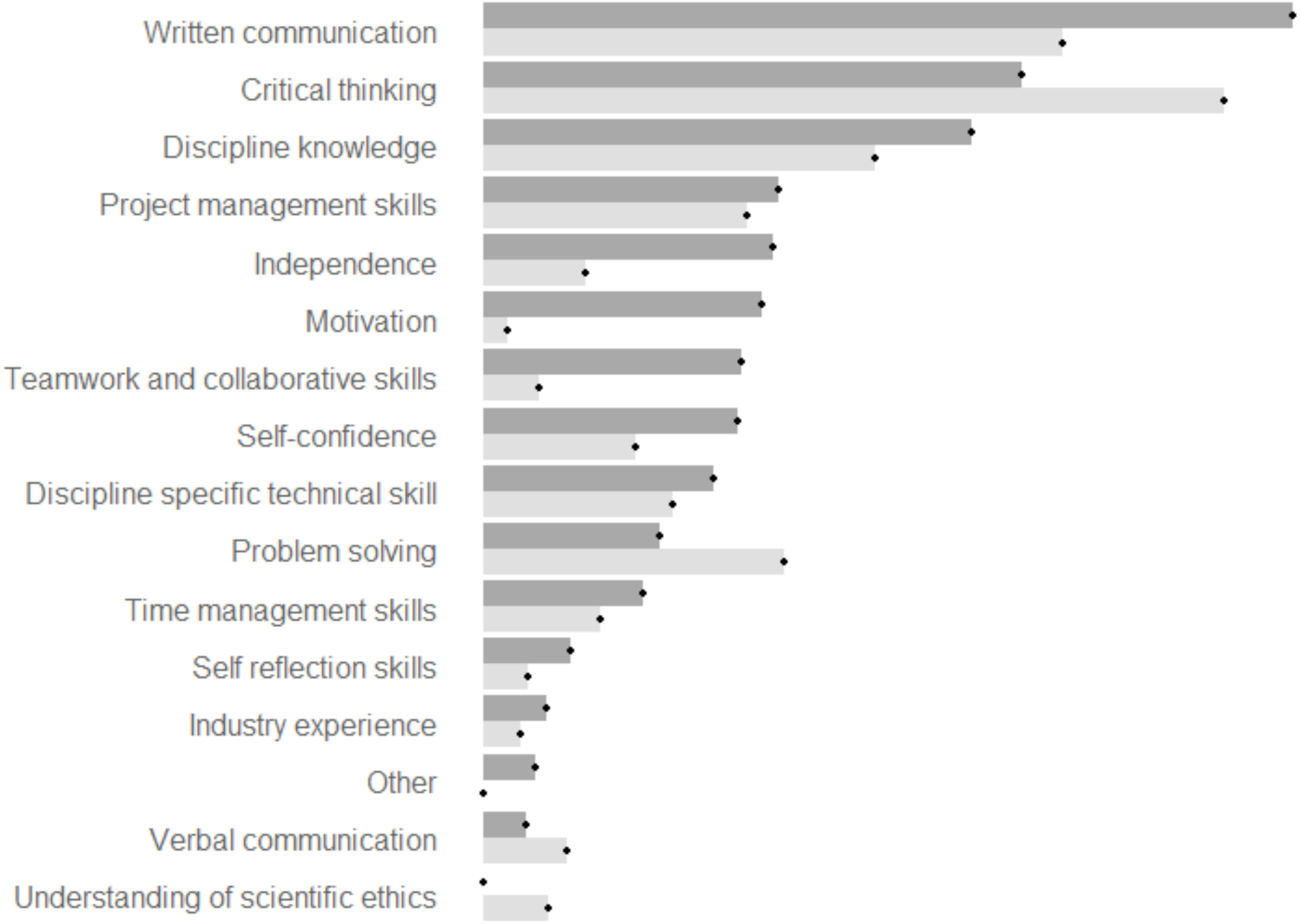
Averaged rank of attributes that supervisors have a responsibility for developing in a candidate as indicated by candidates (dark grey) and supervisors (light grey).

### Guidance given/received

Across attributes, candidates generally considered they were being given less guidance than indicated by the supervisor (Table 3). For written communication, critical thinking, and discipline knowledge (the three attributes identified by the candidates as being the most important to receive supervisor guidance on), candidates considered they got no guidance or only received guidance if they directly asked in 23%, 20% and 29% of cases, respectively, whilst supervisors considered this was almost never the case (2%, 0% and 12% respectively) (Fig. 3). Overall, candidates considered they received guidance at every opportunity or when their supervisor observed they needed it in 64% of cases, whist supervisors perceived they were giving this level of guidance in 82% of cases.

**Table 3.**
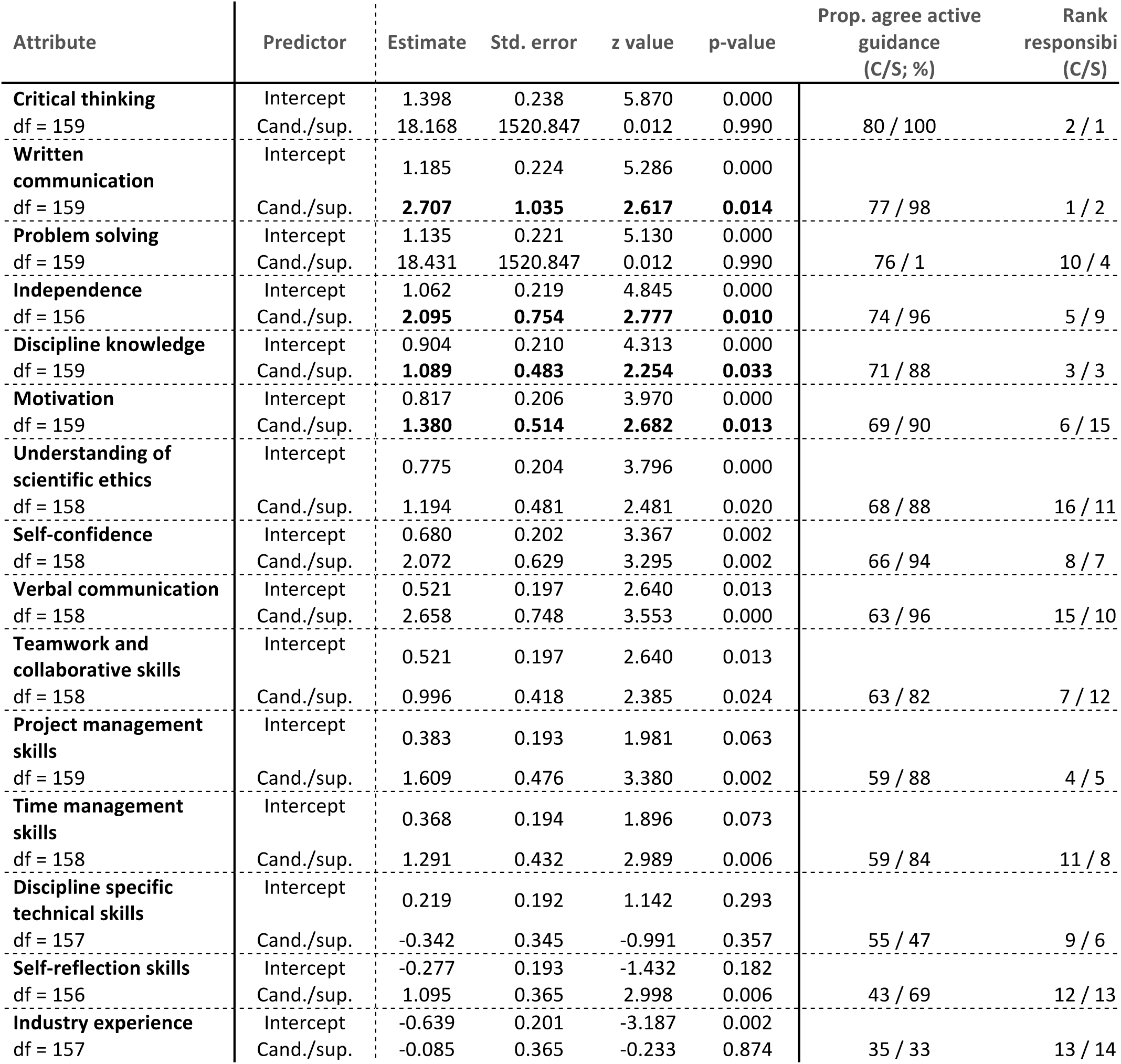
Relationship between perceived levels of guidance received/provided by candidates and supervisor. Positive z values indicate supervisors reporting higher levels of guidance. p-value is FDR corrected. ‘Other’ attribute is not included here and was ranked as C = 14 and S = 16.

**Figure 3.**
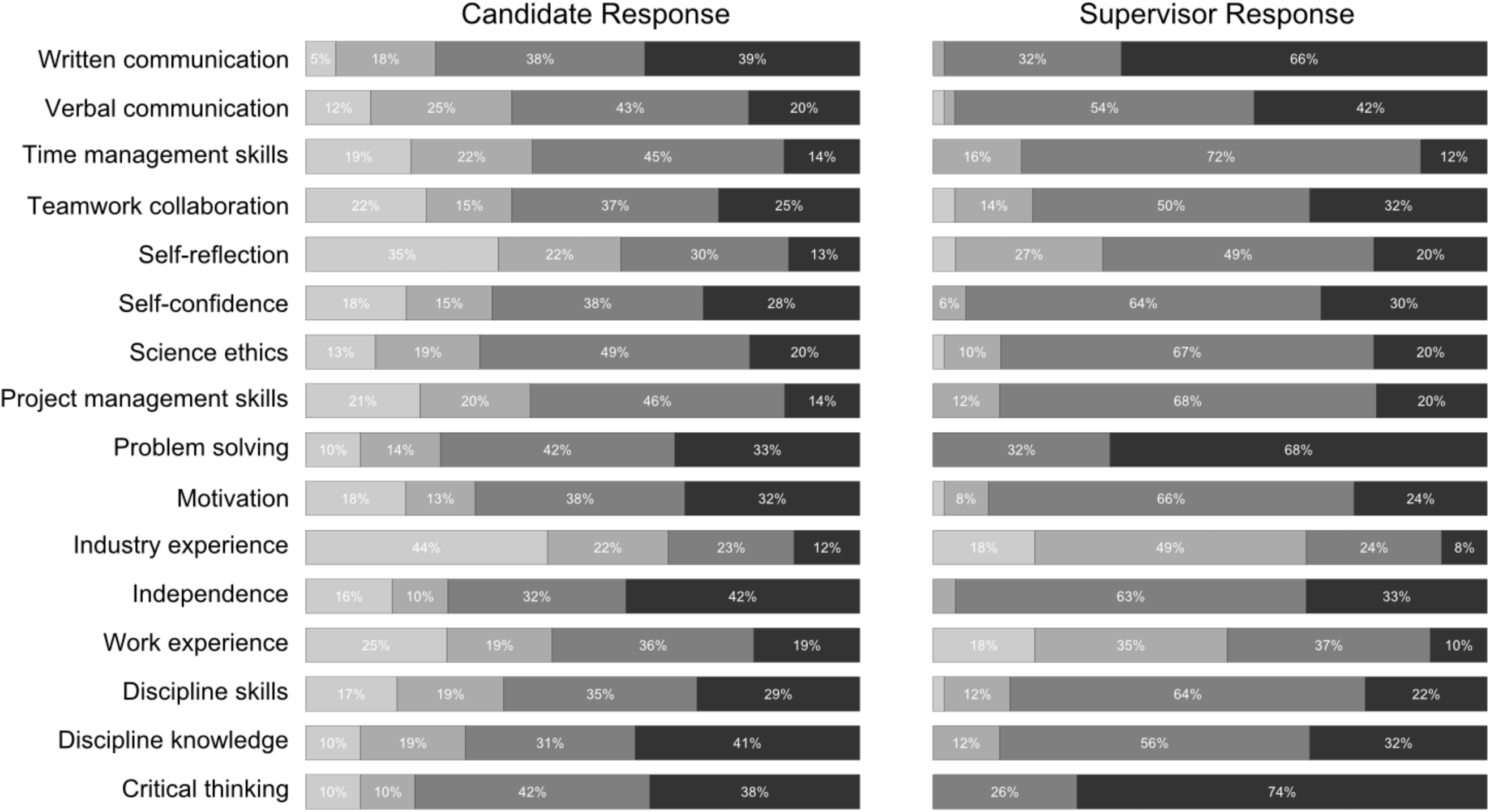
Percentage of respondents who reported levels of guidance received/provided. Guidance levels include *‘none’, ‘only when asked directly’, ‘when I see it is needed’, ‘at every opportunity’* from grey to black respectively.

### Quality of supervision, mental well-being

Candidates reported receiving ‘high’ (34.5%) or ‘very high’ (38.1%) quality PhD supervision, while 7.1% reported receiving ‘very low’ quality supervision, 4% ‘low’ and 15.9% ‘average’. No supervisor reported providing ‘low’ or ‘very low’ quality supervision, 18% reported ‘average’, 68% ‘high’ and 14% ‘very high’.

More than 35% of candidates reported that experiences during their PhD had negatively impacted their overall well-being in a significant way (Fig. 4). Candidates indicated that their mental well-being was significantly influenced in a negative way from personal expectations (54%), research progress (53%), research environment (32%), supervisor expectations (31%) and relationship with supervisor (29%) (Table 4).

**Table 4.**
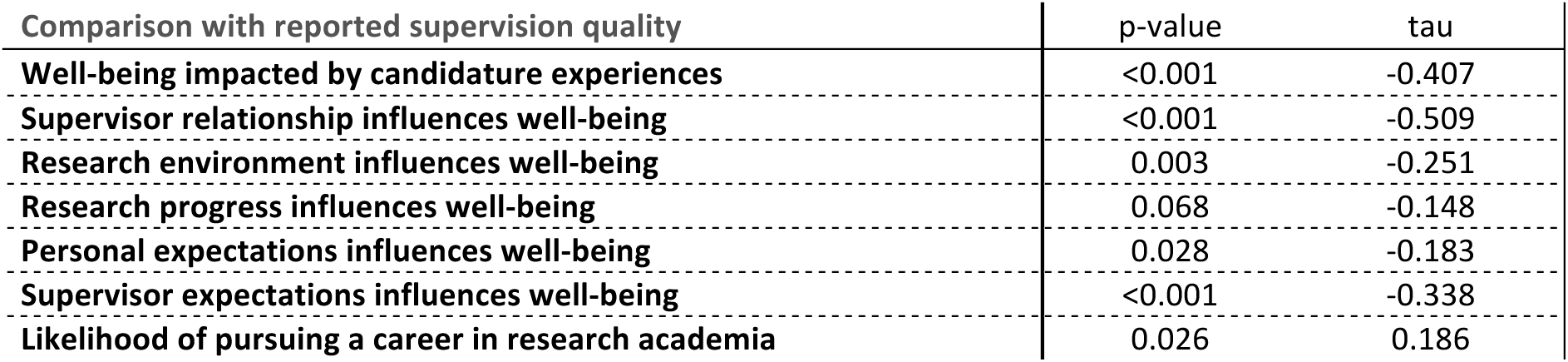
Kendall non-parametric correlation between how candidates reported supervision quality & aspects of well-being, and supervision quality & career aspirations. p-values are FDR corrected.

**Figure 4.**
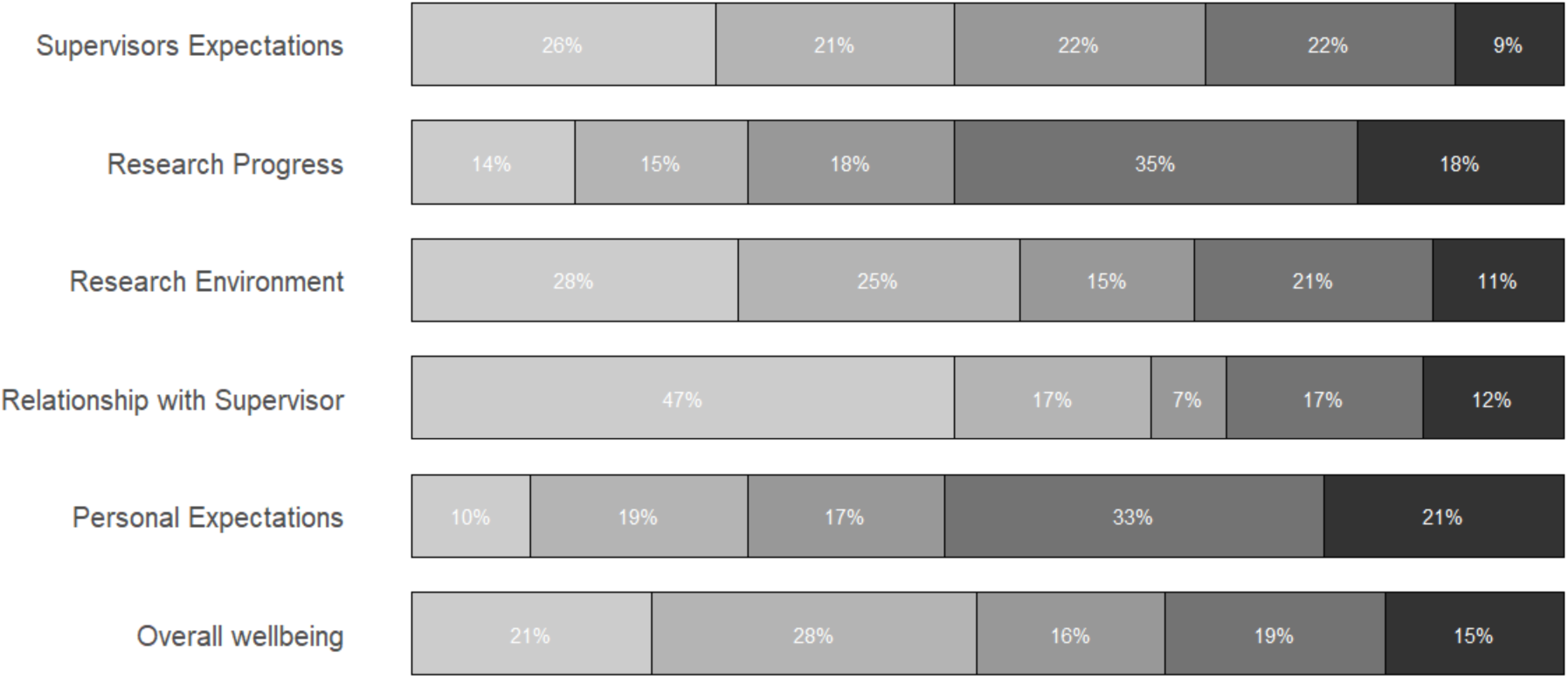
Proportion of candidates indicating whether their PhD experiences have negatively impacted their well-being (overall well-being), and whether specific aspects of candidature has negatively impacted their well-being. From strongly disagreeing that their well-being was negatively impacted on the left (light grey = has not experienced negative well-being) to strongly agreeing on the right (dark grey = has experienced significant negative well-being).

Candidates who reported lower supervision quality were also more likely to report experiencing higher levels of negative well-being, whereas higher quality supervision was related to more positive research career aspirations (Table 4). Female candidates reported receiving higher quality supervision than male candidates (Table 5), yet female candidates reported higher levels of negative well-being compared to male candidates, with supervisor relationship and supervisor expectations being a higher source of negative well-being in particular than reported by males (Table 5).

**Table 5.** Wilcox tests comparing gender differences in experiences of supervisor quality, impacts on negative well-being and career aspirations. p-values are FDR corrected.

| Comparison with gender | p-value | Difference in location | Description |
| --- | --- | --- | --- |
| **Reported supervision quality** | <0.001 | 4 | Female report higher quality supervision |
| **Negative well-being** | <0.001 | 2 | Female report higher levels of negative well-being |
| **Supervisor relationship impacts negative well-being** | <0.001 | 1 | Female report higher levels of negative well-being related to supervisor relationships |
| **Research environment impacts negative well-being** | <0.001 | 2 | Females report bimodal distribution |
| **Research progress impacts negative well-being** | <0.001 | 3 | Males report higher levels of negative well-being related to research progress |
| **Personal expectations impact negative well-being** | <0.001 | 3 | Females skewed towards reporting higher levels, males have bimodal reporting |
| **Supervisor expectations impact negative well-being** | <0.001 | 2 | Females report higher levels of negative well-being related to supervisor expectations |
| **Likelihood of pursuing a career in research academia** | <0.001 | 3 | Female career aspirations are more highly distributed at the end of the options |

## Discussion

In this study we explored how the expectations of PhD candidates and supervisors differ in respect to candidate PhD goals and supervisor guidance, with an aim to promote effective supervisory strategies that increase understanding and collaboration between candidate and supervisor. Our results suggest that the majority of candidates (72.6%) felt that they received better than average supervision quality, and indeed there were many expectations that are in alignment between both parties. For example, at the beginning of candidature, both considered that candidate motivation, enthusiasm for the field and a certain degree of independence are important drivers for candidate success. However, 11.1% of candidates reported receiving lower than average supervision, which is reflected by similar studies in Australian universities (Heath, 2002; McGagh et al., 2016). This dissatisfaction may be driven by important differences in expectations that were observed across respondents. In particular, supervisors strongly considered that candidates should demonstrate good critical thinking skills from the start of candidature. However, candidates considered this less important, instead believing that good academic grades were a more significant demonstration of their ability to do a PhD, despite little evidence for this (Bair & Haworth, 2004). This may lead to conflict if supervisors expect candidates to demonstrate critical thinking when candidates are unsure of what this entails (for example, critical thinking may be confused with ‘book smart’), or do not realise that this is what their supervisor expects from them. In this case, reflection by the supervisor regarding: *i*) whether this is a realistic or fair expectation to have for a new candidate (Ellis et al., 2015), *ii*) how best to develop their critical thinking skills (e.g. giving them relevant articles to review and critique), and *iii*) effective communication to ensure that the candidate is aware that this is an important skill to develop, may all be useful at avoiding misunderstandings and conflicts down the line.

### Outcomes at the end of candidature

By the end of candidature, both candidates and supervisors agreed that the candidate should demonstrate good critical thinking skills, a high level of discipline knowledge and excel at written communication. However, candidates had far higher expectations surrounding the number of publications they would achieve, and placed higher importance on winning awards and grants than did supervisors. This may reflect differences in thinking between candidates and supervisors regarding ultimate goals of the PhD candidature. Supervisors may be primarily concerned with building the requisite skill set required for the candidate to be successful in academia or industry (Gilbert et al., 2004), with the assumption that published papers (and grants) will be an important consequence of these skills (p171; Delamont et al., 2004). Conversely, candidates may be more focussed on quantitative outcomes from their PhD that provide a valuable skill set for future jobs (Roach & Sauermann, 2010), even at the expense of qualitative skills. For example, when candidates are given a lot of help in order to publish quickly, and thereby do not fully develop the written communication skills themselves. Although this potential conflict is well known and discussed amongst academics, with some institutions now promoting PhDs by publication to avoid this conflict (Jackson, 2013), such potentially minor differences in thinking can have profound negative consequences for the candidate. At worst, the candidate may perceive that the supervisor is not working in their best interests if they are working towards different goals. Although the complexity of this scenario increases when supervisors have vested interests in candidate success (MacDonald & Williams-Jones, 2009), it is still likely that invested supervisors still conceptually prioritize candidate skill as a necessary mechanism towards publication, rather than see publications as the overall goal of the PhD. As such, supervisors should be explicitly aware of what the candidate wishes to achieve during their PhD, and guide them towards achievable goals that both promote success in their desired career and meet the requirements of the supervisor and the institution. Critically, communicating that both candidate and supervisor are aiming towards the same specific goals will avoid conflicts near the end of candidature.

### Supervisor guidance

Both candidates and supervisors agreed that supervisors have a strong responsibility to give guidance and feedback on critical thinking, written communication, and relevant discipline knowledge. Interestingly, supervisors considered their responsibility to guide critical thinking and problem solving greater than was expected by candidates. Similar to other studies, candidates instead expected more guidance on developing their academic independence, their collaboration skills, and maintaining motivation (Mowbray & Halse, 2010). Yet, supervisors considered they had little or no responsibility in guiding these less ‘academic’ attributes (Craswell, 2007). This may have disproportionally negative effects on the candidate, with studies consistently showing significant emotional costs when independence is low (De Lange et al., 2004; Levecque et al., 2017; Vanroelen et al., 2009), and when there is little social or collaborative integration within an academic group or institution (Gardner, 2009; Golde, 2000; Kovach Clark et al., 2009; Pyhältö et al., 2009), both of which decrease motivation (Gagné & Deci, 2005; Mason, 2012). Critically, attributes such as independence, self-confidence, collaboration skills, and sustained motivation are all crucial attributes for a successful PhD and career (Mowbray & Halse, 2010; Wuchty et al., 2007), and are qualities that most must learn and develop via mentoring, regular interaction with their research group, and careful self-reflection. Actively supporting and guiding the candidate to increase their autonomy and collaborate with others (either within the research group or with outside collaborators) is likely to have beneficial effects on candidate motivation and productivity (Larivière, 2011), whilst helping them develop independent ideas and teamwork skills (Sinclair et al., 2014). Supervisors may therefore increase the chances of candidate satisfaction and success by actively encouraging the candidate to reflect on how they can develop these crucial qualities.

### Candidate mental-well-being

We show that candidature has negatively influenced over a third of PhD candidates’ surveyed mental well-being, this is in line with findings elsewhere (Evans et al., 2018; Levecque et al., 2017). Our findings that personal expectations, research progress, research environment and supervision impact a relatively large proportion of candidates also echoes previously published word (Barry et al., 2018; Evans et al., 2018). Relationship between candidate and supervisor comes up time and again as a significant contributor to negative well-being. Though, as shown here, the dynamics of how that relationship impacts well-being is sometimes complicated. For instance, despite on average female candidates self-reporting higher levels of supervision quality than male candidates, supervisor expectations and relationship had a greater negative impact on their well-being. There are several possibly explanations for this dynamic that warrants further investigation.

### Effective communication and hidden power dynamics

Our results suggest that there is a disparity between how much guidance supervisors think they are giving to the candidate, and how much candidates perceive they are receiving. In particular, candidates felt they received no guidance at all in 20% of cases, whilst this was very rare amongst supervisors (3%). Worryingly, over 20% of candidates report little guidance (none or only if asked) for critical thinking, written communication and discipline knowledge, despite both parties identifying these subjects as important areas for supervisor guidance. This disparity is likely to stem from the fact that supervisors are time deficient and juggle multiple obligations, whilst candidates in contrast often focus almost solely on obligations surrounding their PhD. Consequently, supervisors may feel that they only have time to give guidance if the candidate specifically asks for it or demonstrates that it is needed. This may be perceived negatively by the candidate, who may interpret the situation as the supervisor not meeting their supervisory obligations. One common suggestion to minimise the chances of this conflict is to spend time discussing and outlining the numerous separate responsibilities and expectations of both candidate and supervisor (Moxham et al., 2013). Although time consuming and requiring a great deal of thought, this is an incredibly worthwhile investment on both sides. Moreover, there are a number of strategies that can help candidates self-reflect and be more productive (Kearns, Forbes, et al., 2008; Kearns, Gardiner, et al., 2008), and supervisors should point students towards these resources when they feel they are not able to meet the needs of the candidate.

However, in addition to this, supervisors should be explicitly aware of power dynamics and its consequences for candidate development and engagement (Grant, 2003; Manathunga, 2007). An inherent discrepancy exists in the major assumptions relating to academic supervision: first is the assumption that both parties are autonomous and rational, and therefore on equal terms to debate and discuss the direction of the candidate. The second assumption is that the supervisor is a wise and knowing authority that also plays the role of examiner, and that the student is a willing disciple in need of guidance (Grant, 2003; Johnson et al., 2000; Manathunga, 2007). These assumptions are inherently in conflict with each other, because in the first scenario both parties are equals, whilst in the second a distinct power imbalance exists. This discrepancy is likely to lead to conflicts when supervisors expect candidates (as rational equals) to bring up problems they may be experiencing, whilst candidates are likely to be submissive or deferential to the authority of the supervisor, and may feel that disagreeing or asking for help is inappropriate or will cause conflict. As such, supervisors should be aware that power dynamics play an important role in mediating candidate behaviour, and that this may discourage candidates from actively pursuing autonomy or, conversely, asking for help if required (e.g. Diamandis, 2017). Although it is challenging for supervisors to walk the line between giving too little guidance (so-called *laissez-faire* supervision) and too much (autocratic supervision)(Delamont et al., 1998; Deuchar, 2008; Gardner, 2008; Van Vugt et al., 2004), awareness that candidates may also be conflicted between dutifully following advice/orders and demonstrating academic independence may help resolve the root of such conflicts when they arise. This conflict may be avoided by consistently applying a democratic leadership style (or ‘participative’ leadership), whereby candidates are encouraged to take a more participative role in the decision making process from the start of candidature. A democratic leadership style has been consistently shown to be the most effective style of leadership to increase performance and satisfaction amongst group members (Eagly et al., 2003; Foels et al., 2000; Van Vugt et al., 2004), and allows the candidate to slowly grow their confidence in their own autonomy within a supportive framework.

### Summary

This study attempts to identify conflicting expectations between PhD candidates and supervisors that are likely to lead to conflict, allowing supervisors to focus efforts on key areas to encourage a successful working relationship. Our results can be summarised by three suggestions to supervisors that may act to reduce conflict and promote positive outcomes for both candidate and supervisor:

1. **Spend time early on in candidature to discuss the importance of, and align each other’s expectations**. For example, supervisors should assist students in developing critical thinking skills from the outset, as our results suggest that candidates are not aware of, or do not place the same level of importance on it as supervisors do early on in their candidature.
2. **Supervisors and candidates should agree to achievable goals that they work towards**. These should include both qualitative (skill sets the candidate should learn) and quantitative (number of papers or grants to be won) outcome. Although these do not need to be overly specific and may evolve over time, this communicates to the candidate that they are both working towards the same set of goals.
3. **Supervisors should play a stronger role in guiding the development of candidate academic independence and collaboration skills**. Both are critical to a successful PhD and career. Supervisors may find that broadening the scope of their supervisory role to actively guide the candidate in developing these qualities will help the candidate maintain motivation and satisfaction over the course of their PhD, and lead to more productive and collaborative research by the candidate.
4. **Maintain effective communication and dialogue throughout**. Supervisors and students should agree on a communication style that best fits both their needs, and regularly evaluate and discuss their communications effectiveness.

We found that supervisors considered that they give more guidance than candidates perceived they receive, and that supervisors often only provided guidance when the candidate asks. We suggest that candidates and supervisors explicitly communicate their separate responsibilities and expectations regarding the spectrum of skills needed to successfully completely a PhD. In addition, supervisors must be cognisant of inherent power dynamics in the student-supervisor relationship, in order to understand and remediate common misalignment of expectations/goals that could lead to dissatisfaction and potential conflict. Applying a democratic leadership style from the outset of candidature may help decrease the effects of this power imbalance.

## References

A. I. McLeod. (2011). Kendall: Kendall rank correlation and Mann-Kendall trend test. https://CRAN.R-project.org/package=Kendall

Bair, C. R., & Haworth, J. G. (2004). Doctoral student attrition and persistence: A meta-synthesis of research. In Higher education: Handbook of theory and research (pp. 481–534). Springer.

Barry, K. M., Woods, M., Warnecke, E., Stirling, C., & Martin, A. (2018). Psychological health of doctoral candidates, study-related challenges and perceived performance. Higher Education Research & Development, 37(3), 468–483. https://doi.org/10.1080/07294360.2018.1425979

Benjamini, Y., & Hochberg, Y. (1995). Controlling the False Discovery Rate: A Practical and Powerful Approach to Multiple Testing. Journal of the Royal Statistical Society. Series B (Methodological), 57(1), 289–300. JSTOR.

Buckley, P. J., & Hooley1, G. J. (1988). The non-completion of doctoral research in management: symptoms, causes and cures. Educational Research, 30(2), 110–120.

Craswell, G. (2007). Deconstructing the skills training debate in doctoral education. Higher Education Research & Development, 26(4), 377–391.

De Lange, A. H., Taris, T. W., Kompier, M. A. J., Houtman, I. L. D., & Bongers, P. M. (2004). Work characteristics and psychological well-being. Testing normal, reversed and reciprocal relationships within the 4-wave SMASH study.

Delamont, S., Atkinson, P., & Parry, O. (2004). Supervising the doctorate. McGraw-Hill Education (UK).

Delamont, S., Parry, O., & Atkinson, P. (1998). Creating a delicate balance: the doctoral supervisor’s dilemmas. Teaching in Higher Education, 3(2), 157–172.

Deuchar, R. (2008). Facilitator, director or critical friend?: Contradiction and congruence in doctoral supervision styles. Teaching in Higher Education, 13(4), 489–500.

Diamandis, E. (2017). A growing phobia. Nature, 544(7648), 129–129.

Eagly, A. H., Johannesen-Schmidt, M. C., & Van Engen, M. L. (2003). Transformational, transactional, and laissez-faire leadership styles: a meta-analysis comparing women and men. American Psychological Association.

Ellis, A. M., Bauer, T. N., Mansfield, L. R., Erdogan, B., Truxillo, D. M., & Simon, L. S. (2015). Navigating Uncharted Waters Newcomer Socialization Through the Lens of Stress Theory. Journal of Management, 41(1), 203–235.

Evans, T. M., Bira, L., Gastelum, J. B., Weiss, L. T., & Vanderford, N. L. (2018). Evidence for a mental health crisis in graduate education. Nature Biotechnology, 36(3), 282–284. https://doi.org/10.1038/nbt.4089

Foels, R., Driskell, J. E., Mullen, B., & Salas, E. (2000). The effects of democratic leadership on group member satisfaction an integration. Small Group Research, 31(6), 676–701.

Gagné, M., & Deci, E. L. (2005). Self-determination theory and work motivation. Journal of Organizational Behavior, 26(4), 331–362.

Gardner, S. K. (2008). “What’s too much and what’s too little?”: The process of becoming an independent researcher in doctoral education. The Journal of Higher Education, 79(3), 326–350.

Gardner, S. K. (2009). Conceptualizing success in doctoral education: Perspectives of faculty in seven disciplines. The Review of Higher Education, 32(3), 383–406.

Gilbert, R., Balatti, J., Turner, P., & Whitehouse, H. (2004). The generic skills debate in research higher degrees. Higher Education Research & Development, 23(3), 375–388.

Golde, C. M. (2000). Should I stay or should I go? Student descriptions of the doctoral attrition process. The Review of Higher Education, 23(2), 199–227.

Grant, B. (2003). Mapping the pleasures and risks of supervision. Discourse: Studies in the Cultural Politics of Education, 24(2), 175–190.

Gu, J., Lin, Y., Vogel, D., & Tian, W. (2011). What are the major impact factors on research performance of young doctorate holders in science in China: a USTC survey. Higher Education, 62(4), 483–502.

Halse, C. (2011). ‘Becoming a supervisor’: the impact of doctoral supervision on supervisors’ learning. Studies in Higher Education, 36(5), 557–570.

Halse, C., & Bansel, P. (2012). The learning alliance: ethics in doctoral supervision. Oxford Review of Education, 38(4), 377–392.

Heath, T. (2002). A quantitative analysis of PhD students’ views of supervision. Higher Education Research & Development, 21(1), 41–53.

Holbrook, A., Shaw, K., Scevak, J., Bourke, S., Cantwell, R., & Budd, J. (2014). PhD candidate expectations: Exploring mismatch with experience. International Journal of Doctoral Studies, 9, 329–346.

Ives, G., & Rowley, G. (2005). Supervisor selection or allocation and continuity of supervision: Ph. D. students’ progress and outcomes. Studies in Higher Education, 30(5), 535–555.

Jackson, D. (2013). Completing a PhD by publication: A review of Australian policy and implications for practice. Higher Education Research & Development, 32(3), 355–368.

Jiranek, V. (2010). Potential predictors of timely completion among dissertation research students at an Australian faculty of sciences. International Journal of Doctoral Studies, 5(1), 1–13.

Johnson, L., Lee, A., & Green, B. (2000). The PhD and the autonomous self: Gender, rationality and postgraduate pedagogy. Studies in Higher Education, 25(2), 135–147.

Kearns, H., Forbes, A., Gardiner, M., & Marshall, K. (2008). When a High Distinction Isn’t Good Enough: A Review of Perfectionism and Self-Handicapping. Australian Educational Researcher, 35(3), 21–36.

Kearns, H., Gardiner, M., & Marshall, K. (2008). Innovation in PhD completion: The hardy shall succeed (and be happy!). Higher Education Research & Development, 27(1), 77–89.

Kiley, M. (2011). Developments in research supervisor training: causes and responses. Studies in Higher Education, 36(5), 585–599.

Kovach Clark, H., Murdock, N. L., & Koetting, K. (2009). Predicting Burnout and Career Choice Satisfaction in Counseling Psychology Graduate Students. The Counseling Psychologist, 37(4), 580–606. https://doi.org/10.1177/0011000008319985

Larivière, V. (2011). On the shoulders of students? The contribution of PhD students to the advancement of knowledge. Scientometrics, 90(2), 463–481.

Levecque, K., Anseel, F., De Beuckelaer, A., Van der Heyden, J., & Gisle, L. (2017). Work organization and mental health problems in PhD students. Research Policy, 46(4), 868–879. https://doi.org/10.1016/j.respol.2017.02.008

Lindén, J., Ohlin, M., & Brodin, E. M. (2013). Mentorship, supervision and learning experience in PhD education. Studies in Higher Education, 38(5), 639–662.

Lovitts, B. E., & Nelson, C. (2000). The hidden crisis in graduate education: Attrition from Ph. D. programs. Academe, 86(6), 44.

MacDonald, C., & Williams-Jones, B. (2009). Supervisor–student relations: Examining the spectrum of conflicts of interest in bioscience laboratories. Accountability in Research, 16(2), 106–126.

Manathunga, C. (2007). Supervision as mentoring: The role of power and boundary crossing. Studies in Continuing Education, 29(2), 207–221.

Mason, M. M. (2012). Motivation, Satisfaction, and Innate Psychological Needs. International Journal of Doctoral Studies, 7, 259–277. https://doi.org/10.28945/1596

McCormack, C. (2004). Tensions between student and institutional conceptions of postgraduate research. Studies in Higher Education, 29(3), 319–334.

McGagh, J., Marsh, H., Western, M., Thomas, P., Hastings, A., Mihailova, M., & Wenham, M. (2016). Review of Australia’s Research Training System. Report for the Australian Council of Learned Academies, http://www.acola.org.au.

Mowbray, S., & Halse, C. (2010). The purpose of the PhD: Theorising the skills acquired by students. Higher Education Research & Development, 29(6), 653–664.

Moxham, L., Dwyer, T., & Reid-Searl, K. (2013). Articulating expectations for PhD candidature upon commencement: ensuring supervisor/student ‘best fit’. Journal of Higher Education Policy and Management, 35(4), 345–354.

Peluso, D. L., Carleton, R. N., & Asmundson, G. J. (2011). Depression symptoms in Canadian psychology graduate students: do research productivity, funding, and the academic advisory relationship play a role? Canadian Journal of Behavioural Science/Revue Canadienne Des Sciences Du Comportement, 43(2), 119.

Pyhältö, K., Stubb, J., & Lonka, K. (2009). Developing scholarly communities as learning environments for doctoral students. International Journal for Academic Development, 14(3), 221–232.

R Core Team. (2017). R: A language and environment for statistical computing. R Foundation for Statistical Computing. https://www.R-project.org/

Roach, M., & Sauermann, H. (2010). A taste for science? PhD scientists’ academic orientation and self-selection into research careers in industry. Research Policy, 39(3), 422–434.

Sinclair, J., Barnacle, R., & Cuthbert, D. (2014). How the doctorate contributes to the formation of active researchers: What the research tells us. Studies in Higher Education, 39(10), 1972–1986.

Tompkins, K. A., Brecht, K., Tucker, B., Neander, L. L., & Swift, J. K. (2016). Who matters most? The contribution of faculty, student-peers, and outside support in predicting graduate student satisfaction. Training and Education in Professional Psychology, 10(2), 102.

Van Vugt, M., Jepson, S. F., Hart, C. M., & De Cremer, D. (2004). Autocratic leadership in social dilemmas: A threat to group stability. Journal of Experimental Social Psychology, 40(1), 1–13.

Vanroelen, C., Levecque, K., & Louckx, F. (2009). Psychosocial working conditions and self-reported health in a representative sample of wage-earners: A test of the different hypotheses of the demand–control–support–model. International Archives of Occupational and Environmental Health, 82(3), 329–342.

Vilkinas, T. (2008). An exploratory study of the supervision of Ph. D./research students’ theses. Innovative Higher Education, 32(5), 297–311.

Wuchty, S., Jones, B. F., & Uzzi, B. (2007). The increasing dominance of teams in production of knowledge. Science, 316(5827), 1036–1039.

