## Supplemental Material for "Supervising the PhD: identifying common mismatches in expectations between candidate and supervisor to improve research training outcomes"

### Supplementary File

#### PhD Candidature Expectations Survey

##### All participants

- Which category do you fall in to? Questions differ for supervisors and candidates, you must answer this question to progress to the correct section.
  - PhD supervisor
  - PhD candidate
  - Recently graduated PhD (<2 yr)
  - Discontinued candidate

##### PhD supervisor questions

- Which gender do you identify as?
  - Female
  - Male
  - Other
  - Prefer not to answer
- What is your age in years? Please answer in numbers not words.
  - [Text input option]
- What is your position title?
  - Associate Lecturer
  - Lecturer
  - Senior Lecturer
  - Associate Professor
  - Professor
  - Other (please specify)
    - [Text input option]
- What is your field of research?
  - Biology
  - Chemistry
  - Physics
  - Information Technology
  - Engineering
  - Mathematics
  - Environmental Science
  - Health/Medicine
  - Other (please specify)
    - [Text input option]
- In what year did you complete your PhD candidature or highest level of education?
  - [Text input option]

**NOTE**: Please answer the following questions in relation to your role as **primary supervisor**.

- How many PhD candidates have you supervised to completion?
  - 0-3

- 4-9
- 10+

- Which top 5 attributes do you expect in a new PhD candidate? Please rank them in order of importance to you.
    - Previous publication(s)
    - Previous awards
    - Good academic grades
    - Project management skills
    - Industry experience
    - Self-confidence
    - Good written communication
    - Good verbal communication
    - Motivated
    - Independent
    - Critical thinking
    - Problem solving
    - Understanding of scientific ethics
    - Cultural fit
    - Reasonable level of discipline knowledge
    - Reasonable level of discipline specific technical skills
    - Enthusiasm for their field
    - Good time management skills
    - Ability to self reflect
    - Demonstrated teamwork skills
    - Other
        - [Text input option]
- Which top 5 attributes or outcomes do you expect a PhD candidate to have achieved by the time of thesis submission? Please rank them in order of important to you.
    - At least 1 publication
    - At least 2 publications
    - At least 4 publications
    - Received award/grants
    - Previous publication/s
    - Previous awards
    - Good academic grades
    - Project management skills
    - Industry experience
    - Self-confidence
    - Good written communication
    - Good verbal communication
    - Motivated
    - Independent
    - Critical thinking
    - Problem solving
    - Understanding of scientific ethics
    - Cultural fit
    - High level of discipline knowledge
    - High level of discipline specific technical skills

    - Enthusiasm for their field
    - Good time management skills
    - Ability to self reflect
    - Teamwork and collaborative skills
    - Other
        - [Text input option]
- Which three attributes do you have the most responsibility for developing in a PhD candidate? Please rank them in order of importance
    - Self-confidence
    - Written communication
    - Verbal communication
    - Motivated
    - Independent
    - Critical thinking
    - Problem solving
    - Understanding of scientific ethics
    - Discipline knowledge
    - Discipline specific technical skills
    - Time management skills
    - Project management skills
    - Industry experience
    - Self reflection skills
    - Teamwork and collaborative skills
    - Other
        - [Text input option]
- What level of guidance do you give to help develop the following attributes in your PhD candidates?
    - Self-confidence
    - Written communication
    - Verbal communication
    - Motivated
    - Independent
    - Critical thinking
    - Problem solving
    - Understanding of scientific ethics
    - Discipline knowledge
    - Discipline specific technical skills
    - Time management skills
    - Project management skills
    - Industry experience
    - Self reflection skills
    - Teamwork and collaborative skills
    - Other
        - [Text input option]
- How many HDR candidates have started a PhD with you but did not submit a PhD thesis? (this includes students that started a PhD but submitted a MPhil)
- In you opinion, rank the quality of the PhD supervision you provide.
    - Very low

- Low
- Average
- High
- Very high

#### Current PhD dandidate questions (and recently graduated or discontinued)

- Which gender do you identify as?
    - Female
    - Male
    - Other
    - Prefer not to answer
- What is your age in years? Please answer in numbers not words.
    - [Text input option]
- What mode of study have you undertaken during candidature.
    - Full-time
    - Part-time
    - Full-time and part-time
- What is/was your field of research?
    - Biology
    - Chemistry
    - Physics
    - Information Technology
    - Engineering
    - Mathematics
    - Environmental Science
    - Health/Medicine
    - Other (please specify)
        - [Text input option]
- In what year did you begin your PhD candidature?
    - [Text input option]

**NOTE**: Please answer the following questions in relation to your **primary supervisor**.

- How many PhD candidates has/had your supervisor previously supervised to completion? (optional)
    - 0-3
    - 4-9
    - 10+
    - Don’t know
- What were you top 5 attributes when you started you candidature? Please rank them in order of importance to you.
    - Previous publication(s)
    - Previous awards
    - Good academic grades
    - Project management skills
    - Industry experience

    - Self-confidence
    - Good written communication
    - Good verbal communication
    - Motivated
    - Independent
    - Critical thinking
    - Problem solving
    - Understanding of scientific ethics
    - Cultural fit
    - Reasonable level of discipline knowledge
    - Reasonable level of discipline specific technical skills
    - Enthusiasm for their field
    - Good time management skills
    - Ability to self reflect
    - Demonstrated teamwork skills
    - Other
        - [Text input option]
- Which top 5 attributes or outcomes do/did you expect to achieve at the time of thesis submission? Please rank them in order of important to you.
    - At least 1 publication
    - At least 2 publications
    - At least 4 publications
    - Received award/grants
    - Previous publication/s
    - Previous awards
    - Good academic grades
    - Project management skills
    - Industry experience
    - Self-confidence
    - Good written communication
    - Good verbal communication
    - Motivated
    - Independent
    - Critical thinking
    - Problem solving
    - Understanding of scientific ethics
    - Cultural fit
    - High level of discipline knowledge
    - High level of discipline specific technical skills
    - Enthusiasm for their field
    - Good time management skills
    - Ability to self reflect
    - Teamwork and collaborative skills
    - Other
        - [Text input option]
- Which three qualities does/did your supervisor have the most responsibility for helping you develop during your PhD candidature? Please rank them in order of importance
    - Self-confidence

    - Written communication
    - Verbal communication
    - Motivated
    - Independent
    - Critical thinking
    - Problem solving
    - Understanding of scientific ethics
    - Discipline knowledge
    - Discipline specific technical skills
    - Time management skills
    - Project management skills
    - Industry experience
    - Self reflection skills
    - Teamwork and collaborative skills
    - Other
        - [Text input option]
- What level of guidance does/did your supervisor provide to help you develop the following attributes during your PhD candidature?
    - Self-confidence
    - Written communication
    - Verbal communication
    - Motivated
    - Independent
    - Critical thinking
    - Problem solving
    - Understanding of scientific ethics
    - Discipline knowledge
    - Discipline specific technical skills
    - Time management skills
    - Project management skills
    - Industry experience
    - Self reflection skills
    - Teamwork and collaborative skills
    - Other
        - [Text input option]
- Experiences during my PhD candidature have negatively affected my mental well-being. (optional)
    - Strongly disagree
    - Disagree
    - Neither agree nor disagree
    - Agree
    - Strongly agree
- Indicate how significantly each of the following negatively influenced your mental wellbeing during your PhD candidature? (optional) [options: Not at all significantly, Not significantly, Neutral, Significantly, Very significantly]
    - Supervisor relationship
    - Research environment
    - Research progress

    - Personal expectations
    - Supervisor expectations
- Considering your PhD experience, how likely are you to pursue a career in research academia?
    - Very unlikely
    - Unlikely
    - Not sure
    - Likely
    - Very likely
- Please rank the quality of the PhD supervision you received.
    - Very low
    - Low
    - Average
    - High
    - Very high
